# Persistence of antibiotic resistance plasmids in bacterial biofilms

**DOI:** 10.1101/098368

**Authors:** Benjamin J. Ridenhour, Genevieve A. Metzger, Michael France, Karol Gliniewicz, Jack Millstein, Larry J. Forney, Eva M. Top

**Affiliations:** Department of Biological Sciences, Institute for Bioinformatics and Evolutionary Studies (IBEST), University of Idaho, Moscow, Idaho, USA; Bioinformatics and Computational Biology Program

**Author notes:** co-first authors (Benjamin J. Ridenhour, Genevieve A. Metzger). Corresponding author: 875 Perimeter Dr, MS 3051, Moscow, ID 83844.

**Keywords:** multiple drug resistance, biofilms, *Acinetobacter baumannii*, plasmids, horizontal gene transfer

## Abstract

The emergence and spread of antibiotic resistance is a crisis in health care today. Antibiotic resistance is often horizontally transferred to susceptible bacteria by means of multi-drug resistance plasmids that may or may not persist in the absence of antibiotics. Because bacterial pathogens often grow as biofilms, there is a need to better understand the evolution of plasmid persistence in these environments. Here we compared the evolution of plasmid persistence in the pathogen *Acinetobacter baumannii* when grown under antibiotic selection in biofilms versus well-mixed liquid cultures. After four weeks, clones in which the plasmid was more stably maintained in the absence of antibiotic selection were present in both populations. On average plasmid persistence increased more in liquid batch cultures, but variation in the degree of persistence was greater among biofilm-derived clones. The results of this study show for the first time that the persistence of MDR plasmids improves in biofilms.

## INTRODUCTION

The emergence and spread of antibiotic resistant bacteria is a crisis faced by healthcare today, and the factors influencing these processes are poorly understood (Centers for Disease Control and Prevention 2013). Resistance to antibiotics can be obtained either through mutations in key genes or by the acquisition of resistance genes via horizontal gene transfer (Benveniste and Davies 1973), which is often mediated by transmissible plasmids (Mazel and Davies 1999; Mathers et al. 2011; Mollenkopf et al. 2016). Broad-host-range plasmids that carry multiple antibiotic resistance genes (MDR plasmids) are of great medical concern because they can be transferred to a wide range of bacterial species (Frost et al. 2005; De Gelder et al. 2007; Mollenkopf et al. 2016). Consequently, a single plasmid transfer event can turn a drug sensitive bacterium into a multiply drug resistant strain. A current alarming example is the rapid worldwide plasmid-mediated spread of antibiotic resistance to colistin, an antibiotic of last resort (Liu et al. 2016; Poirel et al. 2016).

When entering a new bacterial strain, MDR plasmids are often not stably maintained (De Gelder et al. 2007). This instability is overcome by exposure to antibiotics, which impose a strong selective pressure for mutations in the plasmid, host or both, that rapidly improve plasmid persistence (Bouma and Lenski 1988; Heuer et al. 2007; De Gelder et al. 2008; Sota et al. 2010; Millan et al. 2014; Harrison et al. 2015; Loftie-Eaton et al. 2016). We use the term ‘plasmid persistence’, but this trait is also often referred to as plasmid maintenance, plasmid stability, or plasmid retention. The rapid evolution of MDR plasmid persistence has been studied in well-mixed liquid batch cultures (e.g., Bouma and Lenski 1988; Heuer et al. 2007; De Gelder et al. 2008; Sota et al. 2010; Millan et al. 2014; Harrison et al. 2015; Loftie-Eaton et al. 2016), but such culture conditions are not typical in clinical settings where bacteria more naturally occur in biofilms. Until this study, there has been a critical gap in our knowledge of plasmid evolution in biofilms. It is expected that plasmid evolution in biofilms will be different than in liquid cultures because bacterial evolution in general exhibits notably different patterns in biofilms (Rainey and Travisano 1998; Lewis 2001; Donlan 2002; Boles et al. 2004; Ponciano et al. 2009). This is due to the fundamental difference between liquid batch cultures and biofilms, were cells are fixed in space and therefore only compete with neighboring cells. Moreover, the environment a cell experiences (e.g. nutrient availability or antibiotic exposure) is locally variable, which can lead to spatially heterogeneous natural selection and ecology. Thus, some cells may experience particularly strong selection for antibiotic resistance while others may not, simply owing to their location within the biofilm. To combat the spread of MDR plasmids, we need to gain insight into the evolution of bacteria with MDR plasmids in biofilms.

To better understand how biofilm growth affects the evolution of MDR plasmid persistence we performed an experimental evolution study using the Gram-negative pathogen *Acinetobacter baumannii.* Infections caused by *A. baumannii* are an emerging healthcare threat because the organism readily becomes resistant to multiple antibiotics and even pan-drug resistant strains have been reported (Villers 1998; Hsueh et al. 2002; Perez et al. 2007). As a result this species has emerged as an important cause of nosocomial infections. Several studies document longer hospital stays and more severe outcomes for patients with *A. baumannii* infections than with many other bacterial pathogens (Jerassy et al. 2006; Sunenshine et al. 2007). Here we compare the evolution of plasmid persistence in *A. baumannii* when grown in biofilms and well-mixed liquid batch cultures by comparing the persistence of plasmids in evolved clones derived using these two culture conditions. We expected that clones that stably maintain an MDR plasmid would emerge under both conditions. More specifically, by quantitatively analyzing plasmid persistence in individual clones from each population we were able to test two hypotheses. First, weakened selective pressures experienced by bacteria in biofilms will lead to lower average levels of increased plasmid persistence. Second, a broader diversity of plasmid persistence phenotypes will be maintained in biofilms than in liquid cultures, due to the inherent spatial structure of biofilms (Donlan 2002; Boles et al. 2004). Our findings support both of these hypotheses.

## METHODS

### Bacteria and plasmid

The experimental evolution of plasmid persistence in well-mixed liquid cultures and biofilms was done using *Acinetobacter baumannii* ATCC 17978, which was obtained from the American Type Culture Collection (Rockville, MD). This strain of *A. baumannii* is sensitive to tetracycline so plasmid-bearing cells could readily be obtained by plating cells on tetracycline containing media. From here on we refer to it simply as *A. baumannii.*

For this study we used the well-characterized IncP-1 plasmid pB10 (Schlüter et al. 2003) because it has a broad host-range and encodes resistance to four antibiotics (tetracycline, streptomycin, amoxicillin and sulfonamide). The ancestral strain used in all the experiments described here was constructed by electroporation of pB10 into *A. baumannii* (see Appendix A).

### Culture media and conditions

For the evolution experiments *A. baumannii* (pB10) was grown in mineral basal medium (MBM) of M9 salts (Sambrook and Russell 2001) and water supplemented with 18.5 mM succinate, 2 g/L casamino acids, and 10 μg/ml tetracycline (tet), and trace element and mineral mixtures (Wolin et al. 1963) which is hereafter referred to as MBMS-tet. Plasmid persistence assays were done in the same MBMS medium without antibiotics. Dilution plating to obtain individual clones was done using Luria Bertani (LB) agar supplemented with 10 μg/ml tet (LB-tet). All experiments were done at 37°C.

### Experimental evolution protocol

*A. baumannii* (pB10) was evolved in parallel in both biofilms and liquid batch cultures. A time line of the evolution experiments is outlined in Fig. 1. To initiate the experiment, aliquots of an overnight culture grown in MBMS-tet were used to inoculate nine biofilm flow cells that contained ~13 ml of the same medium. These were incubated for 24 h as batch cultures before the flow of fresh medium (5.2 ml/h) was initiated. After four days of growth, which was designated time 0 (t_0_), three flow cells were harvested as follows. The flow cells were moved to a biosafety cabinet where the seal between the lid and body of the flow cells were broken using sterile scalpel blades and the lids were put aside. Media supernatants were removed from the exposed flow cells using a pipette. The biofilms were then resuspended by adding 1mL of 0.85% saline solution and repeatedly pipetting up and down. The resulting suspensions were then transferred to 2mL microcentrifuge tubes and vortexed for 1 minute on high to disperse the cells. Each cell suspension was then examined by microscopy to verify that the cells were dispersed and only free floating cells and small clumps of cells were present. These three cell suspensions were serially diluted and plated on LB-tet agar. After incubation six clones were randomly chosen from each population and archived at −70°C. This entire biofilm harvesting procedure was repeated on days 14 *(t_14_)* and 28 *(t_28_).* Because t_14_ revealed little additional information, we only report on time points t_0_ and t_28_ for the sake of brevity.

**Fig. 1.**
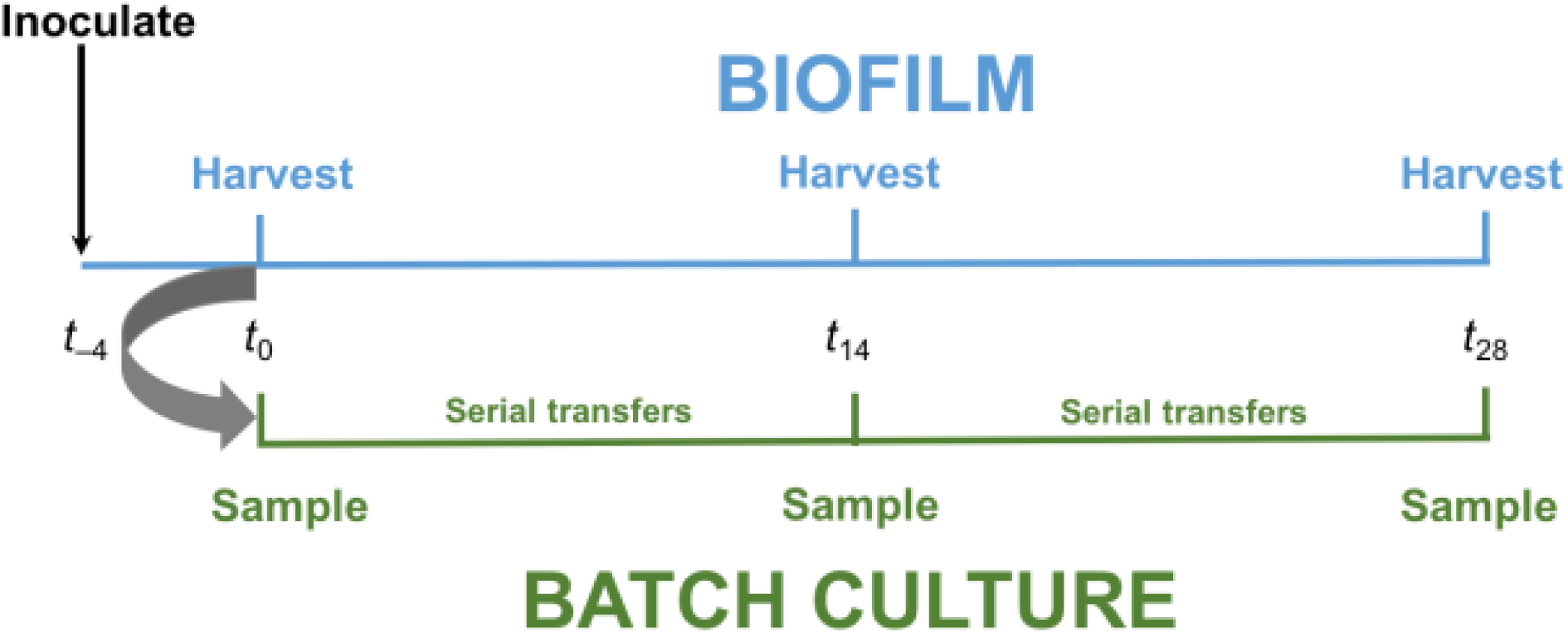
Schematic of experimental evolution experiments in liquid batch cultures (green) and biofilm cultures (blue). Biofilm flow cells were inoculated using an overnight grown culture of the ancestor (*t*_−4_). Four days later (*t*_0_), three randomly selected flow cells were harvested. The resulting cell suspensions were used to inoculate triplicate liquid batch cultures. They were also diluted and plated on LBA-tet to isolate plasmid-bearing clones. The remainder of the cell suspensions were archived at −70°C. The liquid batch cultures were transferred to new media daily for 28 days. On days 14 (*t*_14_) and 28 (*t*_28_), triplicate biofilm cultures were harvested, and samples were taken from the liquid batch cultures.

The liquid batch cultures of *A. baumannii* (pB10) were started by inoculating three replicate test tubes containing five ml of MBMS-tet with 4.9 ul of the cell suspensions from the three biofilms that were harvested at t_0_. These cultures were continuously mixed on rotary shakers at 200 RPM, and each day 4.9 μL of each culture was transferred to 5 mL of fresh media, yielding approximately 10 generations of growth per day. On days 14 and 28, aliquots of these cultures were serially diluted and plated on LB-tet, and six clones from each population were obtained as described above and archived at −70°C.

### Plasmid Persistence Assays

The persistence of plasmids in clonal populations was quantified and compared as follows. Triplicate tubes containing 5 ml of MBMS (without antibiotic) were inoculated with 4.9 μL of each archived clone (representing day 0 of the plasmid persistence assays), and grown overnight. For the next 8 days these cultures were serially transferred to fresh media (4.9 μL into 5mL). On days 0, 5, and 8, a 200 μL sample of each culture was removed, centrifuged, and stored at −20°C. Total DNA was isolated from each culture using a QIAsymphony DSP DNA Mini Kit on a QIAsymphony SP platform (QIAGEN, Inc.). DNA yields were measured fluorometrically using a PicoGreen dsDNA kit.

The fraction of plasmid-bearing cells in each culture was estimated via quantitative PCR (qPCR; Loftie-Eaton et al. 2014) of the plasmid encoded *trfA* gene (encoding the replication initiation protein) and the chromosomally encoded 16S rRNA genes. Plasmid pB10 has a low copy number (~ 2 per cell, data not shown), and the number of 16S rRNA gene copies in *Acinetobacter baumannii* is 5 (Maslunka et al. 2006; Stoddard et al. 2015). The ratio of these two genes was therefore used as a proxy for the fraction of plasmid-bearing cells in the populations, and was expressed as a plasmid:chromosome ratio. These qPCR assays were done in triplicate using a StepOnePlus real-time PCR system and a Power SYBR^TM^ Green PCR master mix (Applied Biosystems Inc.), following the manufacturer’s instructions. Details of the protocol are described in Appendix A.

Analysis of the qPCR data occurred in two stages. The first stage encompassed analysis of the raw qPCR data (i.e. raw fluorescence values for a given PCR cycle in a reaction) for the three qPCR replicates run for each plasmid persistence assay. Given that three replicate plasmid persistence assays were done per clone, 9 qPCR reactions were done for days 5 and 8 of these assays. Because all samples on day 0 came from the same archived glycerol stock and should be homogenous, there were only three qPCR replicates for this day. Statistical analysis of the raw data is described in detail in Appendix A. In a second stage, the qPCR-based estimates of the plasmid:chromosome ratio were used as a measure of the fraction of plasmid-bearing cells. Tracking this ratio over time allowed us to examine differences in plasmid persistence in our 2 different culturing conditions. The log-linear model we used to estimate the rate of plasmid loss over time is described in Appendix A.

Due to variance in our estimates of plasmid:chromosome ratios from the first stage of the analysis, bootstrapping techniques were used to provide more robust confidence intervals in subsequent analyses. In brief, values for ratios were drawn from normal distributions based on the ratios’ estimated means and variances. These randomized values then served as the dependent variable for our log-linear model of plasmid persistence. Additionally, we utilized a Brown-Forsythe-Levene procedure on the residuals of the log-linear model in order to examine the level of phenotypic diversity in biofilm versus liquid cultures (Brown and Forsythe 1974). Such tests are based on group medians to test for linear trends in variances (function ‘ltrend.test’ of the R package ‘lawstat’). The strength and significance of the linear trend between groups is measured using a correlation statistic (negative correlations indicate downward trends and positive correlations indicate upward trends). For both the log-linear models and the Brown-Forsythe-Levene tests, we used a minimum of 1000 bootstrap replicates to determine the significance of our results.

## RESULTS

*A. baumannii* ATCC 17978 containing the MDR plasmid pB10 was grown in biofilms and liquid serial batch cultures for four weeks in the presence of antibiotics selecting for the plasmid (Fig. 1). Plasmid pB10 was shown to be highly unstable in the ancestral strain of *A. baumannii;* the fraction of plasmid bearing cells in the ancestor at the end of persistence assays was only 9 × 10^−5^ (data not shown). Clones isolated from t_0_ biofilms showed levels of instability that were similar to those of the ancestor, and were used as the point of comparison for the evolved host-plasmid pairs isolated from biofilm and liquid batch cultures at *t*_28_.

On average, host-plasmid pairs evolved in both environments for 28 days showed a significant improvement in plasmid persistence when compared to the average level of persistence at t_0_ (Fig. 2). However, the persistence was on average higher for clones evolved in liquid batch cultures than in biofilms, supporting our first hypothesis that natural selection is weaker in biofilms. This was shown in two ways. First, the fraction of plasmid-bearing cells observed in the plasmid persistence assays declined more slowly (Fig. 2, Table 1). Second, the average plasmid bearing fraction at the end of the 8-day plasmid persistence assays was approximately 782 times higher for t28 clones from liquid batch cultures than for the t_0_ clones, while it was only 25 times higher in t28 clones from biofilms (Table 1). To our knowledge, these results show for the first time that plasmid persistence can improve over time in the clinically relevant environment of biofilms, but not to the extent observed in liquid batch cultures.

**Fig. 2.**
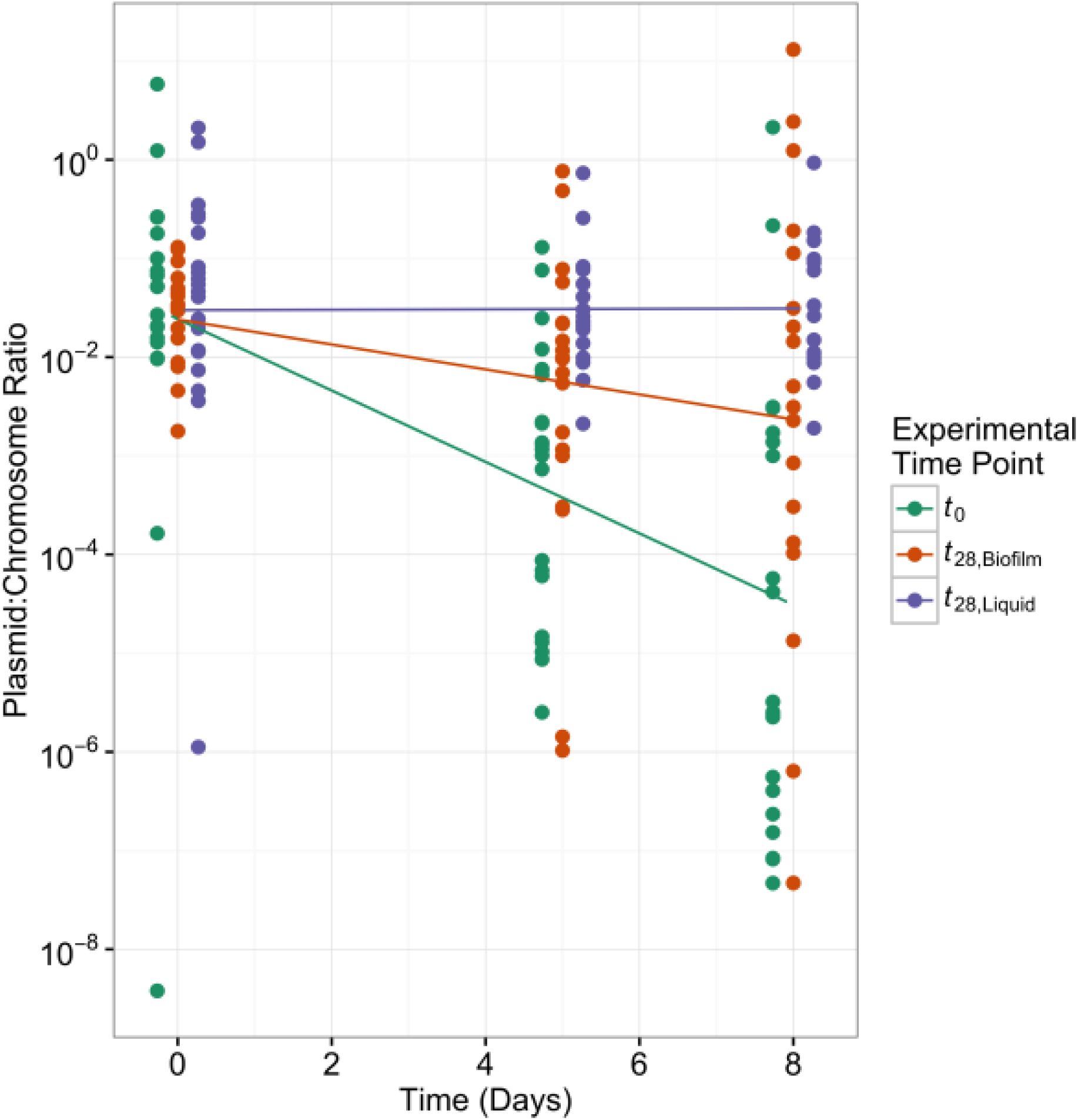
The persistence of plasmid pB10 in clones from liquid batch cultures and biofilm cultures after 28 days of experimental evolution. The fraction of plasmid-bearing cells was determined by quantitative PCR of the plasmid encoded *trfA* gene and chromosomally encoded 16S rRNA genes and expressed as the plasmid:chromosome ratio. The lines show the loss of plasmids over time in populations of clones isolated from biofilms at *t*_0_ (green) and *t*_28_ (red), and liquid batch cultures at *t*_28_ (purple). For each group of samples, the spread of points around their respective lines reflects the diversity of plasmid persistence among the clones from a particular environment.

**Table 1.** Comparison of plasmid loss rates in in clonal populations derived from biofilm and liquid batch cultures. Parameter *β* is the difference in mean rates of decline of the plasmid:chromosome ratio with its associated *p*-value. The fold loss is the relative difference in the fraction of plasmid bearing cells at day 8 of the plasmid persistence assays. The last three columns show differences in diversity between cultures. The first of these (difference in σ_error_) shows the magnitude of the difference in diversity, *ρ* is the test statistic where *H_A_:σ_1_>σ_2_,* and the last column is the *p*-value for *ρ*.

| Comparison | $\beta$ | $p$ | Fold loss | Difference in $\sigma_{\text{error}}$ | $\rho$ | $p$ |
| --- | --- | --- | --- | --- | --- | --- |
| $t_0$ VS. $t_{28}$, liquid | -0.8327 | 0.0008 | 781.8 | 1.6957 | -0.2504 | 0.0097 |
| $t_0$ VS. $t_{28}$, biofilm | -0.4003 | 0.0497 | 24.6 | 0.3612 | -0.0565 | 0.2984 |
| $t_{28}$, biofilm VS. $t_{28}$, liquid | -0.4324 | 0.0320 | 31.8 | 1.3344 | -0.2170 | 0.0367 |

Although plasmid persistence was, on average, higher for clones derived from liquid batch cultures, the variability of plasmid persistence was significantly higher among clones from biofilms (Table 1). These findings provide evidence for our second hypothesis, i.e. that biofilms maintain broader diversity. A visual inspection of the distribution of residuals in the three environments shows that clones from liquid cultures had a much stronger central tendency and less diversity (Fig. 3). A Brown-Forsythe-Levene test indicated that there was no significant difference in the diversity of plasmid persistence when comparing *t*_0_ biofilms and *t*_28_ biofilms. In contrast, the diversity in plasmid persistence in *t*_28_ liquid cultures was significantly lower than in the *t*_0_ cultures used to seed them. Furthermore, we found a significant overall downward trend in diversity from *t*_0_ (most diverse), to *t*_28_ biofilm cultures, to the *t*_28_ liquid cultures (least diverse) (*H_A_:σ_t_0__* > *σ_t_28,biofilm__* > *σ_t_28,liquid__; ρ* = −0.2032; *p* = 0.0120). As hypothesized, these results demonstrate that growth of *A. baumannii* in biofilms maintains higher levels of diversity than growth in liquid cultures.

**Fig. 3.**
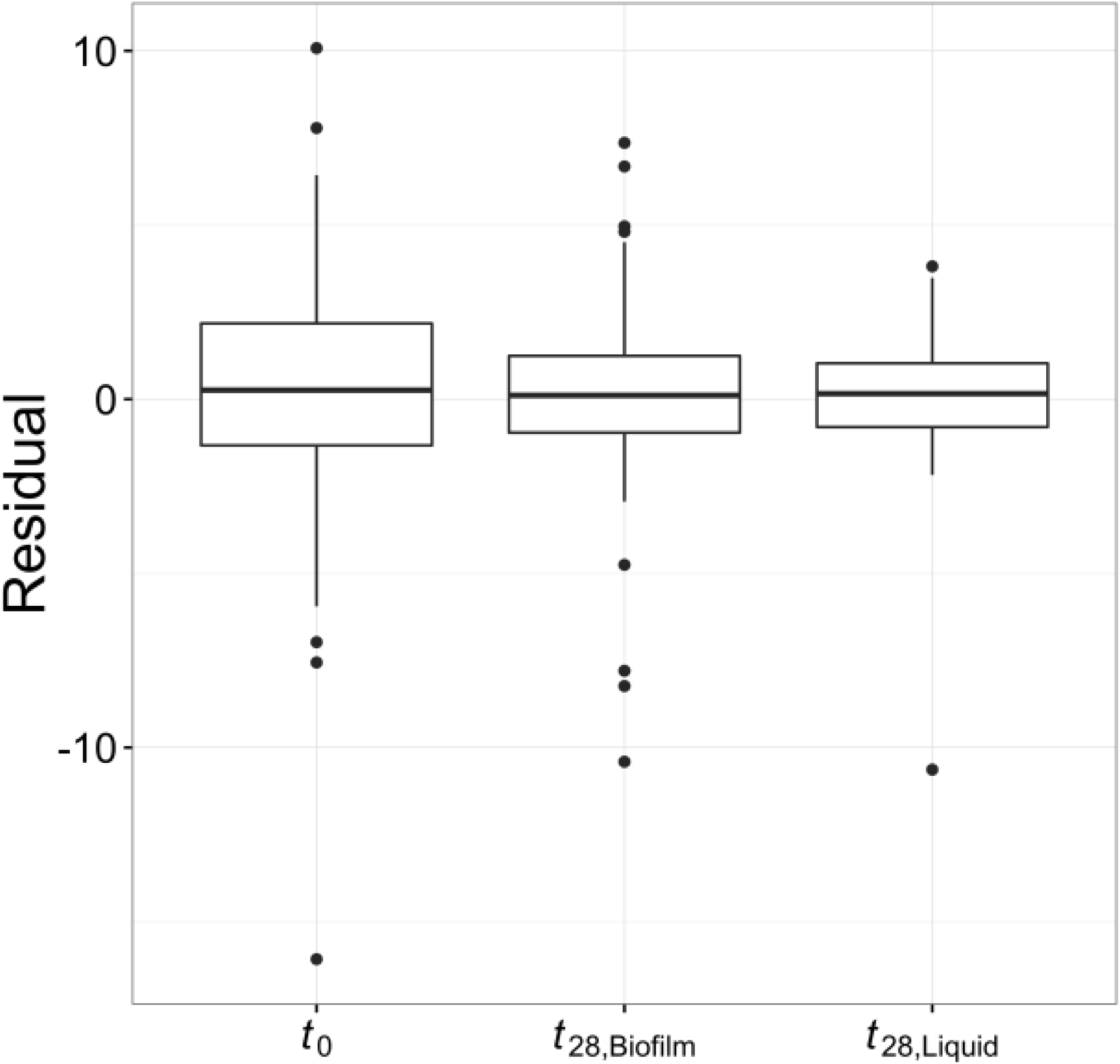
Empirical distributions of residuals within environments. Residuals are the difference between the modeled mean day 8 plasmid:chromosome ratio for a given clone and the modeled mean for a particular environment (e.g. *t*_0_). The variance in residuals represents the phenotypic diversity in a population.

## DISCUSSION

The emergence and spread of antibiotic resistant bacteria is one of the greatest crises facing healthcare today, yet the factors influencing these processes are still poorly understood. Since self-transmissible MDR plasmids play a key role in the spread of antibiotic resistance (Frost et al. 2005), we need to better understand how well they persist in pathogens in the absence of selection and how this persistence may evolve over time in clinically relevant conditions such as biofilms. So far as we know, all former studies addressing plasmid evolution were done using well-mixed liquid cultures (e.g., Bouma and Lenski 1988; Heuer et al. 2007; De Gelder et al. 2008; Sota et al. 2010; Millan et al. 2014; Harrison et al. 2015; Loftie-Eaton et al. 2016). The results of this study show for the first time that the persistence of MDR plasmids improves in biofilms, but less so than in liquid cultures due to a higher phenotypic diversity in biofilms.

There are multiple reasons why the persistence of an MDR plasmid is on average lower and more diverse after four weeks of antibiotic exposure in biofilms than in liquid batch cultures. One of the differences between these two environments is that bacterial growth rates vary widely in biofilms. This is due to the inherent heterogeneity of spatially structured biofilm environments, which leads to gradients of nutrients, electron acceptors and metabolic waste products that govern bacterial metabolism (Boles et al. 2004). Cells at the biofilm surface grow more readily than those buried deep in the biofilm matrix where nutrient limitation hampers cell growth. This could be important because plasmid loss only occurs during cell division when plasmids are not properly apportioned between the daughter cells. Thus, one potential explanation for the lower average improvement and higher diversity in plasmid persistence in biofilms is that MDR plasmids may persist in this environment simply by remaining in non-dividing cells, on which natural selection cannot act.

Another important difference between biofilms and liquid cultures is that cells are fixed in space in biofilms, which limits competitive interactions to a cell’s immediate neighbors. Natural selection therefore occurs on a local scale. This localized scale of selection in biofilms has been shown to result in the accumulation of genetic diversity (Boles et al. 2004; Ponciano et al. 2009). In accordance with this, our large replicated evolution experiment showed that after four weeks biofilm populations harbor a greater diversity of plasmid persistence than well-mixed liquid batch cultures (Fig. 3). This large design allowed us to assess the diversity generated by antibiotic application, rather than simply estimating means. Under the conditions used in this study the variation in plasmid persistence was roughly the same in four day old biofilms (*t_0_*) and those grown for another four weeks in the presence of tetracycline *(t_28_).* In contrast, the diversity at *t_0_* was drastically reduced when the populations were used to found liquid batch cultures that were subsequently grown for four weeks (Table 1, Fig. 3). Our findings confirm that growth in biofilms either protracts or prevents the selective sweeps commonly observed in liquid cultures (Martens and Hallatschek 2011). Thus, biofilms maintain diversity in MDR plasmid persistence that would not have been observed in the traditional experimental evolution studies. This phenomenon can be thought of as a ‘seed bank’ of genetic diversity (Boles et al. 2004), which may reduce the effectiveness of future antibiotic treatments because pre-adapted clones may be present. Therefore, understanding how growth within a biofilm affects the evolution and persistence of MDR plasmids will provide fundamental insights to the emergence and recalcitrance of bacterial infections. We are currently determining the genetic basis of improved plasmid persistence using high-throughput sequencing technologies.

In conclusion, the data presented support a hypothesis that is of medical concern: biofilms generate and maintain a broad diversity of antibiotic resistant bacteria that better retain an MDR plasmid than the ancestor, but with variable success. To combat the emergence of antibiotic resistance we should not only find alternative therapies that limit MDR plasmid persistence in biofilms of pathogens, but investigate whether the genetic diversity maintained in biofilms facilitates outbreaks of MDR pathogens in the future.

## ACKNOWLEDGMENTS

We would like to thank Matthew Settles, Thibault Stalder, Wesley Loftie-Eaton, Silvia Smith, Rachana Regmi, Andrew Avery, Sean West, and Taylor Wilkinson for helpful discussion on the design of this project, collection of data, and comments on previous versions of the manuscript. This project has been funded by DOD-DM110149. GAM was also in part supported by a fellowship from the Institute for Bioinformatics and Evolutionary Studies and the Bioinformatics and Computational Biology graduate program at the University of Idaho.

## Appendix A Experimental and Analytical Details

### Construction of ancestral *Acinetobacter baumannii* host-plasmid pair

To allow the host to adapt to the culture environment prior to experimental evolution with plasmid pB10, the archived *Acinetobacter baumannii* strain ATCC17978 was grown for 10 days with daily passage of 0.1% of the volume in fresh mineral medium (MBM) supplemented with 18.5 mM succinate as the main carbon source and 2 g/L casamino acids, hereafter named MBMS. MBM consists of 1X M9 salts (Sambrook and Russell, 2001), amended with 10 mL of a vitamin stock solution, and 10 mL trace element stock solution per liter (Wolin et al. 1963).

Following the medium adaptation phase, plasmid pB10 was introduced into *A. baumannii* by electroporation with selection on tetracycline (tet, 10 μg/ml). After electroporation 10 colonies were selected and grown overnight in MBMS with tetracycline (10 μg/ml), hereafter called MBMS-tet; plasmid DNA extractions were performed to determine the presence of pB10. Clones that appeared to contain intact pB10 were digested with the restriction enzymes *PstII* and *HindIII* and the patterns were compared to a control of pB10 DNA extracted from *Escherichia coli* to confirm the presence of intact pB10. The presence of full size pB10 was confirmed in one clone, which was then selected as the ancestor for all experiments.

To obtain a large liquid culture that contained as little genetic diversity among cells as possible, an extinction-dilution procedure was performed. A dilution series was made from 10^−1^ to 10^−10^ and used to inoculate large volumes of liquid media. These cultures were then allowed to grow until approximately 8 hours after the cultures inoculated from the 10^−8^ dilution showed turbidity. Because none of the more dilute cultures showed growth in that time period, the 10^−8^ culture was selected as the ancestral stock and 0.65 mL aliquots were archived at −70°C with 30% glycerol.

### Experimental Evolution

Biofilms were grown in flow cells that were set up in a manner similar to that of Ponciano et al. (2009), with some modifications described here and in the main text of this manuscript. The flow cell chambers and top and bottom slides were made of polycarbonate plastic and the chambers were sealed using silicone adhesive. The flow cells were inoculated with 200μL of our archived ancestral culture, initially clamped for 24 hours to allow the cells to settle, and then subsequently unclamped to initiate a flow of MBMS-tet at a rate of 5.4 mL/hr. The flow cells, media, and waste bottles were contained within an incubator that was maintained at 37°C. Media and waste bottles were changed approximately every 2-3 days as needed. All flow cells were checked daily for overgrowth into the tubing supplying the media and for leaks. Leaking flow cells were removed and tubing and filters with substantial overgrowth were replaced as needed.

To harvest the biofilms the top plate of the flow cell was removed with a sterile scalpel blade. The media supernatant was pipetted off and the biofilm population was resuspended in 2mL of phosphate buffer saline (PBS). A portion of the suspension was diluted in PBS and plated onto LB agar supplemented with tet (LB-tet). The remainder of the harvested cell suspension was combined with glycerol and stored in 0.65 mL aliquots in the −70°C freezer.

Four days after inoculation, designated *t_0_*, 4.9μL portions of each of the three biofilm cell suspensions were used to inoculate each of three test tubes containing 5 mL of MBMS-tet (that is, each t0 biofilm was used to inoculate one of the liquid culture replicates); this represented the starting point of the liquid batch cultures, *t_0_*. These liquid cultures were then grown at 37°C in a shaking incubator in serial batch cultures: 4.9μL was transferred into 5 mL of MBMS-tet every 24 ± 1 hours. Of each liquid cultures a 1 mL sample was archived every 5 days, as well as at 14 days (*t_14_*) and 28 days (*t*_28_) after inoculation (*t*_0_). Here we only report on the populations harvested at *t*_28_.

Six colonies were randomly chosen from the LB-tet agar plates for each biofilm or liquid culture harvested at both the *t*_0_ and *t*_28_ time points. They were grown overnight in liquid MBMS-tet before being archived at −70°C with glycerol. The presence of pB10 was confirmed in these clones using a plasmid extraction protocol, followed by restriction enzyme digestion and gel electrophoresis. These clones were then tested for plasmid persistence.

### Plasmid extraction procedure

We used the plasmid extraction procedure described in Selim and Hagag (2013) with the following modifications: 1) the volume of TE, lysis solution, and 2M Tris-HCl was doubled, 2) RNAse was added along with the 100 μL of TE before the lysis step, and 3) centrifugation at 20,000xg and 4 °C was done for 45 minutes after adding Tris-HCl, followed by transferring the supernatant into phenolchloroform.

### DNA purification and qPCR conditions

To purify DNA for qPCR we used the DNA extraction protocol provided with QIAsymphony DSP DNA Mini kits. Briefly, 5 μl of liquid overnight cultures of *A. baumannii* was added to 220 μl of buffer ATL and transferred to 2 ml barcoded micro tubes. Next, 20 μl of proteinase K was added to the samples, followed by a brief vortexing and 1 hr of shaking at 900 rpm in a shaker-incubator (at 56°C). Later, the samples were briefly spun down to remove the condensation from tube caps. Four microliters of RNAse A (100 mg/ml) was then added and followed by a 2 min incubation period at room temperature. All tubes with cellular lysate were then transferred to the sample carrier trays for the QIAsymphony SP instrument set to run a DNA Blood & Tissue LC200 protocol (default parameters) with a 50 μl elution volume.

The amplification parameters for all qPCR reactions were 94°C for 10 min, 40 cycles of 94°C for 15 s, and 60°C for 60 s. Parameters for melting curves were 94°C for 15 s and 60°C for 30 s, followed by a temperature increase to 94°C with 0.1% ramp rate. The fluorescent signal was acquired after each 60°C amplification step and collected continuously during the melting curve analysis. 0.2 ng of template DNA was used per qPCR reaction, with every sample assayed in triplicate. On plasmid pB10 gene *trfA,* encoding the replication initiation protein, was amplified using primers 5’-GAACAGCACCACGATTTCGG-3’ (forward) and 5’-TACTACACAAGGGCCGAGGA-3’ (reverse). The *trfA* copies were compared to the 16S rRNA genes in the *A. baumannii* chromosome, amplified using primers 1080*γ*F (5’-TCGTCAGCTCGTGTYGTGA-3’) and *γ*1202R (5-CGTAAGGGCCATGA TG-3’) described previously (Bachetti De Gregoris et al. 2011).

### Data analysis

Analysis of the raw qPCR fluorescence data was done using a Bayesian hierarchical model. We implemented our model in the Stan programming language and the R statistical programming language (Stan is available in R via the package RStan; Hoffman and Gelman, 2014). Stan utilizes a Hamilontian Monte Carlo (HMC) algorithm to sample posterior distributions of model parameters. Conceptually, the model being used is based on the simple enzyme kinetic model *x* + *y* → 2*x* where *x* is the concentration of template DNA and *y* is concentration of PCR primer. This model has the time dependent solution

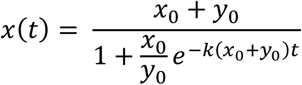

where *x*_0_ is the starting DNA concentration, *y*_0_ is the starting primer concentration, *k* is the rate at which Taq polymerase catalyzes the reaction, and *t* is time measured in PCR cycles. We estimate the quantity x0 because it represents the initial concentration of either *trfA* or 16S rRNA. A hierarchical model allowed us to model the error to account for the fact that the error of replicate retention assays should be correlated with the error of the qPCR replicates, thus resulting in more accurate parameter estimates. The HMC sampler was run for 4 independent chains consisting of 2000 generations, the first 1000 of which were discarded as a burn-in period; further details can be found in the documentation for the Stan language (http://mc-stan.org). Potential scale reduction factor (PSRF) values were checked to verify that convergence was obtained across the 4 chains.

The qPCR-based *trfA/16S* rRNA ratio estimates were used as a measure of the fraction of plasmid-bearing cells. Specifically, we utilized a log-linear model to determine the rate at which plasmids were lost from *A. baumannii* cultures and the variation in that rate. The simple linear model was

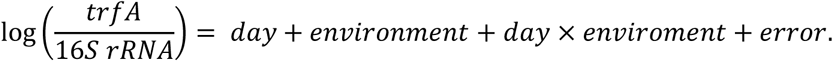

The log transformation of the ratio improved model diagnostics and corrected for heteroscedasticity in residuals. Significant *day × environment* terms indicate differences in the rate of plasmid loss between growth environments. Furthermore, the residual variance for each environment (*σ*_*error*|*environment*_) from this model is an estimate of the biodiversity of plasmid persistence phenotypes.

